# Airway epithelial interferon response to SARS-CoV-2 is inferior to rhinovirus and heterologous rhinovirus infection suppresses SARS-CoV-2 replication

**DOI:** 10.1101/2021.11.20.469409

**Authors:** Elizabeth R. Vanderwall, Kaitlyn A. Barrow, Lucille M. Rich, David F. Read, Cole Trapnell, Oghenemega Okoloko, Steven F. Ziegler, Teal S. Hallstrand, Maria P. White, Jason S. Debley

## Abstract

**Introduction:** Common alphacoronaviruses and human rhinoviruses (HRV) induce type I and III interferon (IFN) responses important to limiting viral replication in the airway epithelium. In contrast, highly pathogenic betacoronaviruses including SARS-CoV-2 may evade or antagonize RNA-induced IFN I/III responses.

**Methods:** In airway epithelial cells (AECs) from children and older adults we compared IFN I/III responses to SARS-CoV-2 and HRV-16, and assessed whether pre-infection with HRV-16, or pretreatment with recombinant IFN-β or IFN-λ, modified SARS-CoV-2 replication. Bronchial AECs from children (ages 6-18 yrs.) and older adults (ages 60-75 yrs.) were differentiated *ex vivo* to generate organotypic cultures. In a biosafety level 3 (BSL-3) facility, cultures were infected with SARS-CoV-2 or HRV-16, and RNA and protein was harvested from cell lysates 96 hrs. following infection and supernatant was collected 48 and 96 hrs. following infection. In additional experiments cultures were pre-infected with HRV-16, or pre-treated with recombinant IFN-β1 or IFN-λ2 before SARS-CoV-2 infection.

**Results:** Despite significant between-donor heterogeneity SARS-CoV-2 replicated 100 times more efficiently than HRV-16. IFNB1, INFL2, and CXCL10 gene expression and protein production following HRV-16 infection was significantly greater than following SARS-CoV-2. IFN gene expression and protein production were inversely correlated with SARS-CoV-2 replication. Treatment of cultures with recombinant IFNβ1 or IFNλ2, or pre-infection of cultures with HRV-16, markedly reduced SARS-CoV-2 replication.

**Discussion:** In addition to marked between-donor heterogeneity in IFN responses and viral replication, SARS-CoV-2 elicits a less robust IFN response in primary AEC cultures than does rhinovirus, and heterologous rhinovirus infection, or treatment with recombinant IFN-β1 or IFN-λ2, markedly reduces SARS-CoV-2 replication.

## INTRODUCTION

The novel coronavirus SARS-CoV-2 has rapidly infected humans across the globe, causing one of the most devastating pandemics in modern history, with over 240 million confirmed cases and nearly 5 million deaths worldwide by October 2021(1). While most cases of the resulting coronavirus disease 2019 (COVID-19) are mild, some cases are severe and complicated by respiratory and multi-organ failure(2), with a fatality rate ranging from as low as 0.2% to as high as 27% depending on underlying medical co-morbidity and age(3). For the first half of the pandemic, incidence of COVID-19 was surprisingly low among children(3), however, there is evidence that SARS-CoV-2 infection rates are as high in children as older adults(3) and that children can shed SARS-CoV-2 while asymptomatic and for prolonged periods(4). More recently, the incidence of COVID-19 in the United States has increased significantly among children and adolescents(5). Understanding mechanisms that explain the heterogeneity of severity with SARS-CoV-2 infection between individuals and across different age groups may assist efforts to develop therapeutic interventions to treat and prevent COVID-19.

One potential explanation for the wide variation in COVID-19 disease severity is the differences in the innate immunity between individuals, particularly the heterogeneity of type I and III interferon (IFN) responses. Innate immune sensing of coronaviruses, including SARS-CoV-2, is thought to occur primarily through pattern recognition receptors (PRRs) including the cytosolic RIG-I-like receptors, melanoma differentiation-associated protein 5 (MDA5; coded for by the gene *IFIH1),* and retinoic acid-inducible gene I (RIG-I) as well as cell surface or endosomal transmembrane toll-like receptors (TLRs) TLR3 and TLR7, which lead to the activation of signaling cascades that further induce type I and III IFN responses(6) (7) (8) (9). Common human coronavirus (HCoV) strains (e.g. alpha-coronavirus strain 229E) potently induce type I and III IFN, and their replication is susceptible to inhibition by IFN I/III, leading to suppression of the early phase of viral replication(10) (11). In contrast, previous highly pathogenic beta-HCoVs (e.g. SARS-CoV and MERS-CoV) encode viral proteins with a greater capability to antagonize RNA-induced type I and III IFN production through perturbation of RNA sensing(12) (13) (14) (15) (16) (17). Likewise, IFN responses at mucosal surfaces appear to be muted during SARS-CoV-2 infection as compared to other respiratory viruses, suggesting evasion of innate immune responses by SARS-CoV-2(18) (19). Data from our lab and others indicates that epithelial infection with human rhinovirus increases the expression of the entry receptors for SARS-CoV-2(20) (21), suggesting that when these two viruses concurrently infect individuals the response to one virus could modulate the response to the other.

Data from clinical studies increasingly support a hypothesis that deficiency of initial IFN responses to SARS-CoV-2 may allow for increased viral replication that then supports systemic inflammatory responses that contribute to COVID-19 pathology and severity(19) (22) (23) (24). Ziegler et al. recently performed scRNA-seq on nasopharyngeal swabs from 15 healthy adults, 14 adults with mild COVID-19 and 21 adults with severe COVID-19, and observed that epithelial cells from patients with severe COVID-19 had less robust expression of anti-viral interferon response genes as compared to patients with mild COVID-19 and healthy controls supporting their conclusion that a “failed” nasal epithelial innate anti-viral response may be a risk factor for severe COVID-19(25).

The objectives of our study were to determine if heterogeneity in bronchial epithelial type I and III IFN responses to SARS-CoV-2 between individual pediatric and adult donors was associated with SARS-CoV-2 replication, to compare airway epithelial IFN responses between SARS-CoV-2 and human rhinovirus-A16 (HRV-16), and to determine the effects of HRV pre-infection or exogenous IFN treatment on SARS-CoV-2 replication in organotypic airway epithelial cell (AEC) cultures from children and adults. We hypothesized that type I and III IFN responses would be less vigorous to SARS-CoV-2 than to HRV infection, that IFN responses would be associated with SARS-CoV-2 replication, and that HRV pre-infection and/or recombinant IFN treatment of airway epithelial cultures would decrease replication of SARS-CoV-2. Some of the results of these studies have been previously reported in the form of an abstract(26).

## METHODS

Bronchial AECs from children ages 6-18 years (n=15) and older adults ages 60-75 years (n=10) were differentiated *ex vivo* at an air-liquid interface (ALI) to generate organotypic cultures. AECs from children were obtained under study #12490 approved by the Seattle Children’s Hospital Institutional Review Board. Parents of subjects provided written consent and children over 7 years of age provided assent. Primary bronchial AECs from adults were purchased from Lonza® or obtained from a tracheal segment lung transplant donor lung tissue. AECs were differentiated *ex vivo* for 21 days at an ALI on 12-well collagen-coated Corning® plates with permeable transwells in PneumaCult™ ALI media (Stemcell™) at 37°C in an atmosphere of 5% CO2 as we have previously described, producing an organotypic differentiated epithelial culture with mucociliary morphology(27) (28) (29) (30).

Experimental conditions in this study included: infection of AECs with SARS-CoV-2 alone, infection of AECs with HRV-16 alone, infection of AECs with HRV-16 followed by infection with SARS-CoV-2 72 hours later, infection of IFNβ1 treated AECs with SARS-CoV-2, and infection of IFNλ2 treated AECs with SARS-CoV-2. For AECs treated with recombinant IFN, recombinant IFNβ1 (1ng/mL) or IFNλ2 (10ng/mL) was added to basolateral transwell chamber with every medium change, starting 72 hours prior to SARS-CoV-2 infection and continuing until 96 hours following SARS-CoV-2 infection. The concentrations of recombinant I FNβ1 and IFNλ2 were chosen based on data from preliminary experiments in three primary AEC lines comparing the effect of a range of concentrations of each cytokine from 0.1 - 10 ng/mL on SARS-CoV-2 replication (data not shown). In a Biosafety Level 3 (BSL-3) facility, cultures were infected with SARS-CoV-2 isolate USA-WA1/2020 or HRV-16 at a multiplicity of infection (MOI) of 0.5. At 96 hrs. following SARS-CoV-2, or following HRV-16 infection alone, RNA was isolated from cells using Trizol® and protein was isolated from cell lysates with RIPA buffer (Sigma-Aldrich®) containing Triton X100 1% and SDS 0.1%, methods that we have demonstrated fully inactivate SARS-CoV-2(31).

Expression of *IFNB1, IFNL2, CXCL10, IFIH1, and GAPDH* were measured by quantitative polymerase chain reaction (qPCR) using Taqman® probes. To measure SARS-CoV-2 replication in AEC cultures we used the Genesig® Coronavirus Strain 2019-nCoV Advanced PCR Kit (Primerdesign®), with duplicate assays of harvested RNA from each SARS-CoV-2-infected AEC experimental condition. The viral copy number used in analyses of each experimental condition was the mean of duplicate assays from each experimental condition. Similarly, to measure HRV-16 replication in AEC cultures we used the Genesig® Human Rhinovirus Subtype 16 PCR Kit (Primerdesign®).

To extract protein from the cell layer of SARS-CoV-2-infected AEC cultures, media was first removed from the basolateral chamber of transwells. Next, 100 μL of cold PBS was added to the apical surface of cultures and 1mL was added to the basolateral chamber of cultures as a wash step. Next, 50μL of RIPA buffer for protein extraction ready-to-use-solution (Sigma-Aldrich®, Product No. R0278) containing Triton X100 1% and SDS 0.1% was added to the apical surface of AECs and incubated for 15 minutes on ice. A pipet tip was then used to gently scratch each apical well in a crosshatch pattern to loosen AECs from the transwell membrane. Material was collected, centrifuged at 10,000 rpm at 4°C for 10 minutes, then supernatant containing isolated protein was collected. IFNβ1, IFNλ2, and CXCL-10 protein concentrations in cell lysates, and IFNβ1, IFNλ3, and CXCL-10 concentrations were measured in cell culture supernatants, via a Human Luminex® Assay (R&D®), with protein concentrations normalized to total protein levels in lysate (BCA assay; Sigma-Aldrich®).

### Statistical Analysis

Gene expression and protein levels are presented as means +/− standard deviation (SD) when data were normally distributed, and as medians with interquartile range if one or more groups were not normally distributed. To determine if data was normally distributed the Kolmogorov-Smirnov test was used (alpha = 0.05). *IFNB1, IFNL2, IFIH1 and CXCL10* relative expression were standardized using *GAPDH* as a non-regulated housekeeping gene. GenEx version 5.0.1 was used to quantify gene expression from qPCR normalized to GAPDH (MultiD Analyses AB, Göteborg, Sweden) based on methods described by Pfaffl(32). Data in at least one group or condition in each experiment analyzed were determined to be non-normally distributed, therefore nonparametric tests were used for analyses. To compare gene expression data and distributions of protein concentrations in cell lysates and supernatants between paired groups the Wilcoxon matched-pairs signed rank test was used. For unpaired data the Mann-Whitney test was used for analyses. For experiments with three or more conditions the Kruskal-Wallis one-way ANOVA on ranks test was used, and *post hoc* comparisons between pairs of subject groups were made using Dunn’s multiple comparisons test (significance level set at p<0.05). Correlations were determined using the Spearman’s rank correlation coefficient. Data was analyzed using Prism® 9.0 software (GraphPad Software Inc., San Diego, CA.). Statistical significance was set at *p*<0.05.

## RESULTS

In organotypic primary bronchial AEC cultures from children (n=15) and older adults (n=10) we observed marked heterogeneity in SARS-CoV-2 replication between human donors (Figure 1). The clinical characteristics of human airway epithelial donors included in these experiments is summarized in Table 1. Despite the significant between-subject heterogeneity in SARS-CoV-2 replication, we observed that SARS-CoV-2 replicated approximately 100 times more efficiently than HRV-16 in these primary bronchial AEC cultures (Figure 1; SARS-CoV-2 median copy number 215,387 vs. HRV-16 median copy number 2211; p<0.0001) when parallel cultures from each donor were infected with each virus at the same MOI of 0.5. When data from pediatric and adult cultures were analyzed separately SARS-CoV-2 replication was also markedly greater than HRV-16 in cultures within each donor age group (children: SARS-CoV-2 median copy number 215,387 vs. HRV-16 median copy number 2602; p<0.001; adults: SARS-CoV-2 median copy number 75,940 vs. HRV-16 median copy number 2184; p=0.002). SARS-CoV-2 replication was not significantly different between AEC cultures from pediatric and adult donors (median copy number 215,387 vs. 75,940; p=0.23), and among pediatric donors SARS-CoV-2 replication was not significantly different between cultures from children with asthma and healthy children (median copy number 60,540 vs. 436,465; p=0.3).

**Figure 1.**
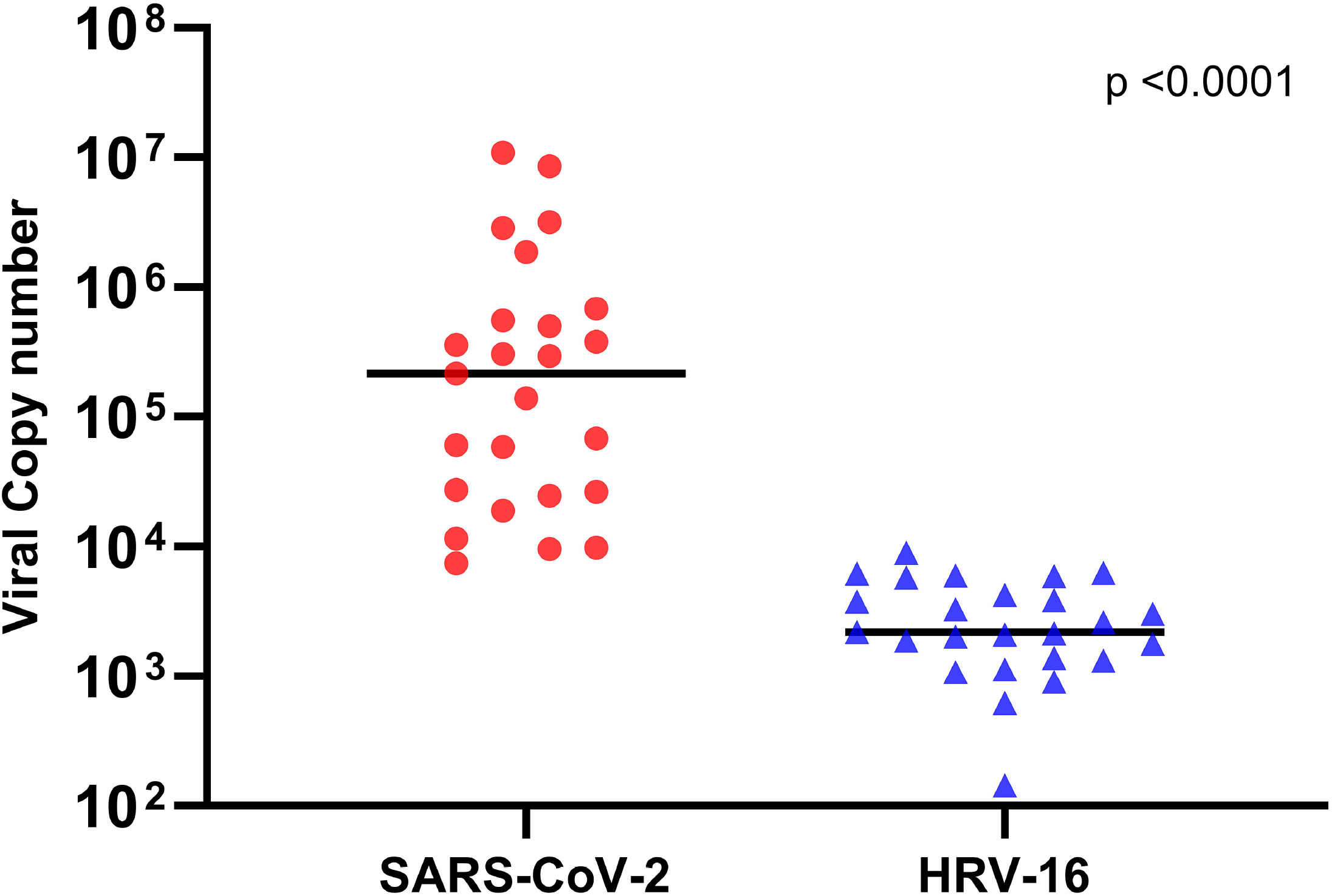
SARS-CoV-2 and HRV-16 replication by quantitative PCR in primary bronchial AECs from children (n=15) and adults (n=10). Viral copy number was quantified by PCR in RNA harvested from AEC cultures 96 hours following infection (MOI of 0.5) with either SARS-CoV-2 (red circles) or HRV-16 (blue triangles). SARS-CoV-2 replication was significantly greater than HRV-16 (median copy number 215,387 vs. HRV-16 median copy number 2211; p<0.0001 by Wilcoxon matched-pairs signed rank test; bars indicate median values).

**Table 1:** Airway Epithelial Cell Donor Characteristics.

|  | **Pediatric AEC Donors**<br>**(n=15)** | **Adult AEC Donors**<br>**(n=10)** |
| --- | --- | --- |
| **Age** (yrs., mean +/- SD) | 10.5 +/- 2.0 | 67 +/- 4.9 |
| **Gender** (female) | 9 (60%) | 4 (40%) |
| **Active Smoker** | 0 (0%) | 3 (30%) |
| **History of Asthma** | 8 (53%) | 0 (0%) |
| **Obesity** | 0 (0%) | 4 (40%) |
| **Hypertension** | 0 (0%) | 3 (3%) |

**AEC = Airway epithelial cell**

For primary bronchial epithelial cultures wherein SARS-CoV-2 and HRV-16 infection was compared in parallel, RNA harvested 96 hours following infection was available from 22 donor cultures (n=14 children, n=8 adults) to allow measurement of *IFNB1, IFNL2,* and *CXCL10* gene expression, and protein was available from cell lysate collected 96 hours following infection from 20 donor cultures (n=12 children, n=8 adults) to allow for measurement of IFNβ1, IFNλ2 (IL-28A), and CXCL-10 protein levels. As compared to uninfected cultures, the relative increase in expression of *IFNB1* following infection with HRV-16 was significantly greater than following infection with SARS-CoV-2 (median increase expression 4.4-fold vs. 1.4-fold, p<0.0001; Figure 2, panel A). Similarly, the relative increase in expression of *IFNL2* following infection with HRV-16 was significantly greater than following infection with SARS-CoV-2 (median increase expression 21.2-fold vs. 4.3-fold, p<0.0001; Figure 2, panel C), as was the increase in expression of *CXCL10* (median increase expression 9.8-fold vs. 5.4-fold, p=0.003; Figure 2, panel E). The expression of these three genes was significantly greater following HRV-16 infection than following SARS-CoV-2 in cultures from both children and adults when analyzed separately (data not shown). The concentrations of IFNβ1, IFNλ2 (IL-28A), and CXCL-10 protein, normalized to total protein concentration, in cell lysates collected 96 hours following infection with HRV-16 were also significantly greater than in parallel cultures following SARS-CoV-2 infection (IFNβ1: median 892 vs. 663 pg/mL, p=0.02, Figure 2, panel B; IFNλ2 (IL-28A): 9848 vs. 7123 pg/mL, p=0.02, Figure 2, panel D; and CXCL-10: 69,306 vs. 15,232 pg/mL, p<0.0001, Figure 2, panel F).

**Figure 2.**
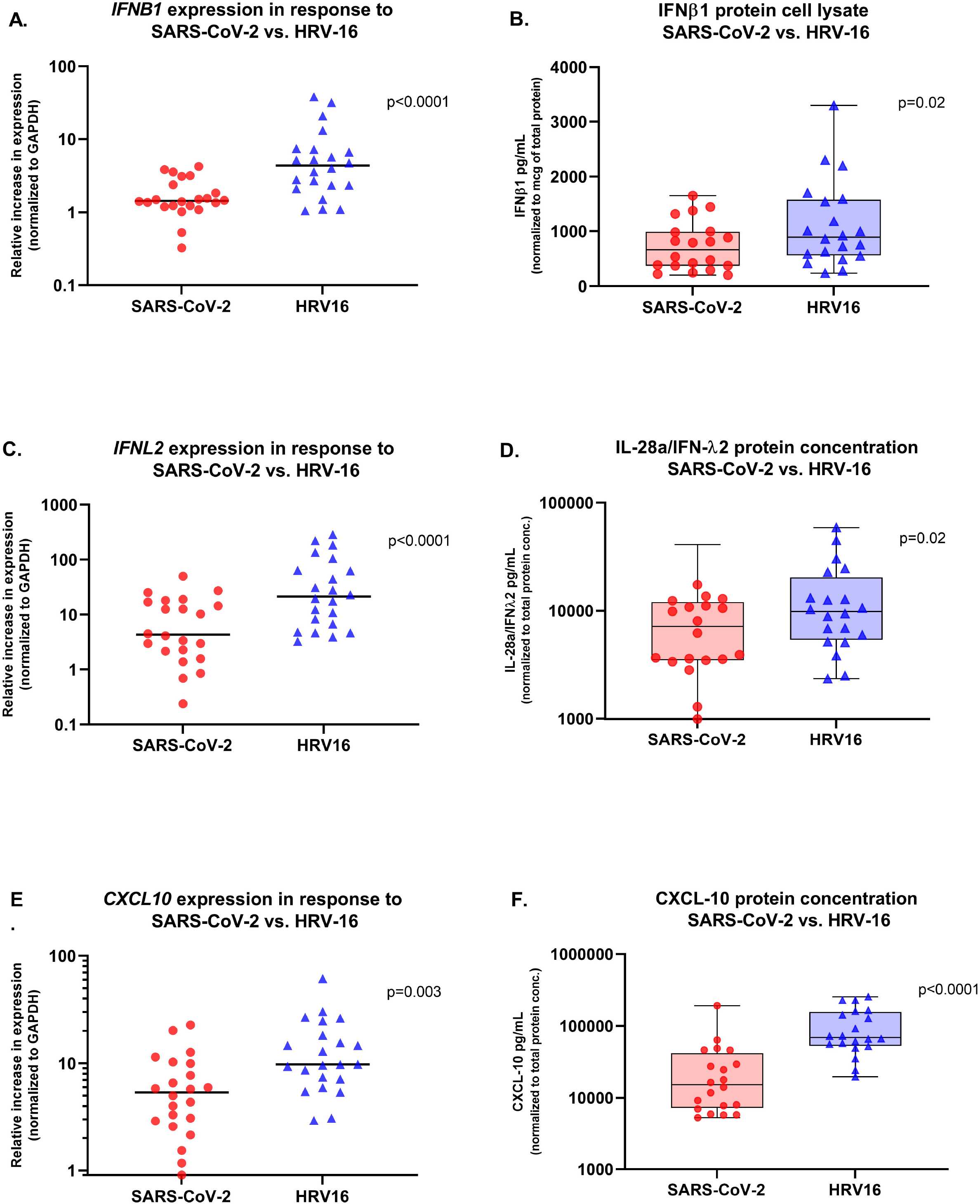
Relative gene expression of *IFNB1, IFNL2,* and *CXCL10* (normalized to GAPDH expression) by primary bronchial airway epithelial cell cultures in children (n=14) and adults (n=8), and parallel IFNβ1, IFN-λ2 (IL-28a), and CXCL10 protein concentrations in cell lysates (normalized to total protein concentration), from primary bronchial airway epithelial cell cultures from children (n=12) and adults (n=8) harvested 96 hours after SARS-CoV-2 (red circles) or HRV-16 (blue triangles) infection. Expression of *IFNB1* and corresponding concentrations of IFNβ1 in cell lysates were significantly greater in cultures after infection with HRV-16 than in cultures infected with SARS-CoV-2 (**Panel A**, median increase expression 4.4-fold vs 1.4-fold, p<0.0001; **Panel B**, median 892 pg/mL vs 663 pg/mL, p=0.02). Expression of *IFNL2,* and IFNλ2 protein concentrations in cell lysates, were significantly greater following HRV-16 infection than SARS-CoV-2 infection (**Panel C**, median increase expression 21.2-fold vs 4.3-fold, p<0.0001; **Panel D**, median 9848 pg/mL vs 7123 pg/mL, p=0.02). Expression of *CXCL10,* and CXCL10 protein concentrations in cell lysates, were significantly greater following HRV-16 infection as compared to SARS-CoV-2 infection (**Panel E**, median increase expression 9.8-fold vs 5.4-fold, p=0.003; **Panel F**, 69,306 pg/mL vs 15,232 pg/mL, p<0.0001). Analyses by Wilcoxon matched-pairs signed rank test. Bars indicate median values. Boxplots indicate interquartile range and whiskers indicate minimum and maximum values.

Of cultures wherein SARS-CoV-2 and HRV-16 infection was compared in parallel, supernatant was collected from n=16 donor cell lines 48 hours following infection and from n=20 cell lines 96 hours following infection. Concentrations of I FNβ1 in supernatant (normalized to total protein concentration) were higher at 48 hours vs. 96 hours post infection for both viruses. However, IFN β1 concentrations were significantly greater following HRV-16 as compared to SARS-CoV-2 infection at both 48 hours (median 60.4 vs. 12.5 pg/mL, p<0.001, Figure 3, panel A) and 96 hours (median 7.1 vs. 1.4 pg/mL, p<0.001, Figure 3, panel A). IFNλ2 (IL-28A) concentrations were below the assay detection level in supernatants for most samples (data not shown). IFNλ3 (IL-28B) concentrations in supernatants were significantly greater following HRV-16 as compared to SARS-CoV-2 infection at both 48 hours (median 1335 vs. 40.6 pg/mL, p<0.001, Figure 3, panel B) and 96 hours (median 197 vs. 48 pg/mL, p<0.001, Figure 3, panel B). CXCL10 concentrations in supernatants were also significantly greater following HRV-16 as compared to SARS-CoV-2 infection at both 48 hours (median 293,805 vs. 10,407 pg/mL, p<0.001, Figure 3, panel C) and 96 hours (median 179,858 vs. 150,939 pg/mL, p=0.04, Figure 3, panel C).

**Figure 3.**
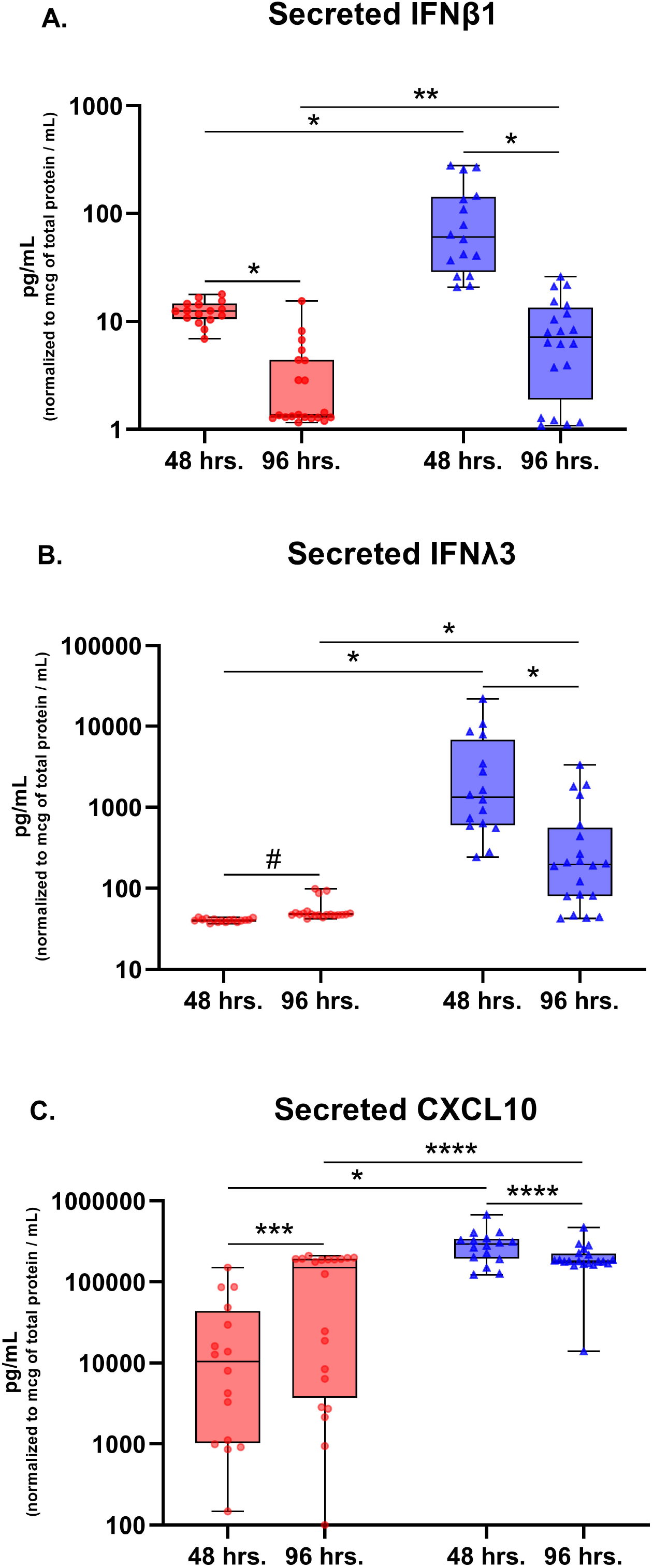
Concentrations of secreted IFNβ1, IFN-λ3, and CXCL10 (normalized to total protein concentration) in the supernatant of primary bronchial epithelial cell cultures 48 hours and 96 hours after infection with SARS-COV-2 (red circles) or HRV-16 (blue triangles). Secreted IFNβ1 concentrations peaked at 48 hours post viral infection (**Panel A**), and IFNβ1 concentrations were significantly higher in HRV-16 infected cultures than SARS-CoV-2 infected cultures at both time points (**Panel A**). IFNλ3 (**Panel B**) and CXCL10 (**Panel C**) concentrations were also significantly greater in HRV-16 infected cultures at 48 and 96 hours following infection. *p<0.001, **p=0.005, ***p=.0.03, ****p=0.04, #p=0.2. Analyses by Mann–Whitney tests. Boxplots indicate interquartile range and whiskers indicate minimum and maximum values.

At 96 hours following infection we assessed correlations between relative expression of *IFNB1* and *IFNL2* in individual primary bronchial epithelial cell lines and viral replication (SARS-CoV-2 copy number) in those cultures. Both *IFNB1* and *IFNL2* gene expression was inversely correlated with SARS-CoV-2 replication *(IFNB1* r=-0.61, p=0.003; *IFNL2* r=-0.42, p=0.05; Figure 4). Because concentrations of IFNβ1 and IFNλ3 in supernatants were highest at 48 hours following infection, we assessed correlations between supernatant concentrations of these cytokines at 48 hours following SARS-CoV-2 infection and viral replication at 96 hours following infection and observed a significant inverse correlation between supernatant IFNβ1 concentrations and viral replication (r=-0.53, p=0.02; Figure 5, panel A) and a trend toward an inverse correlation between supernatant IFNλ3 concentrations and viral replication (r=-0.44, p=0.06, data not shown). We observed significant negative correlations between CXCL10 protein concentrations in both supernatant (r=-0.56, p=0.01, data not shown) and cell lysate (r = - 0.65, p=0.002; Figure 5, panel B) at 96 hours following infection and SARS-CoV-2 replication.

**Figure 4.**
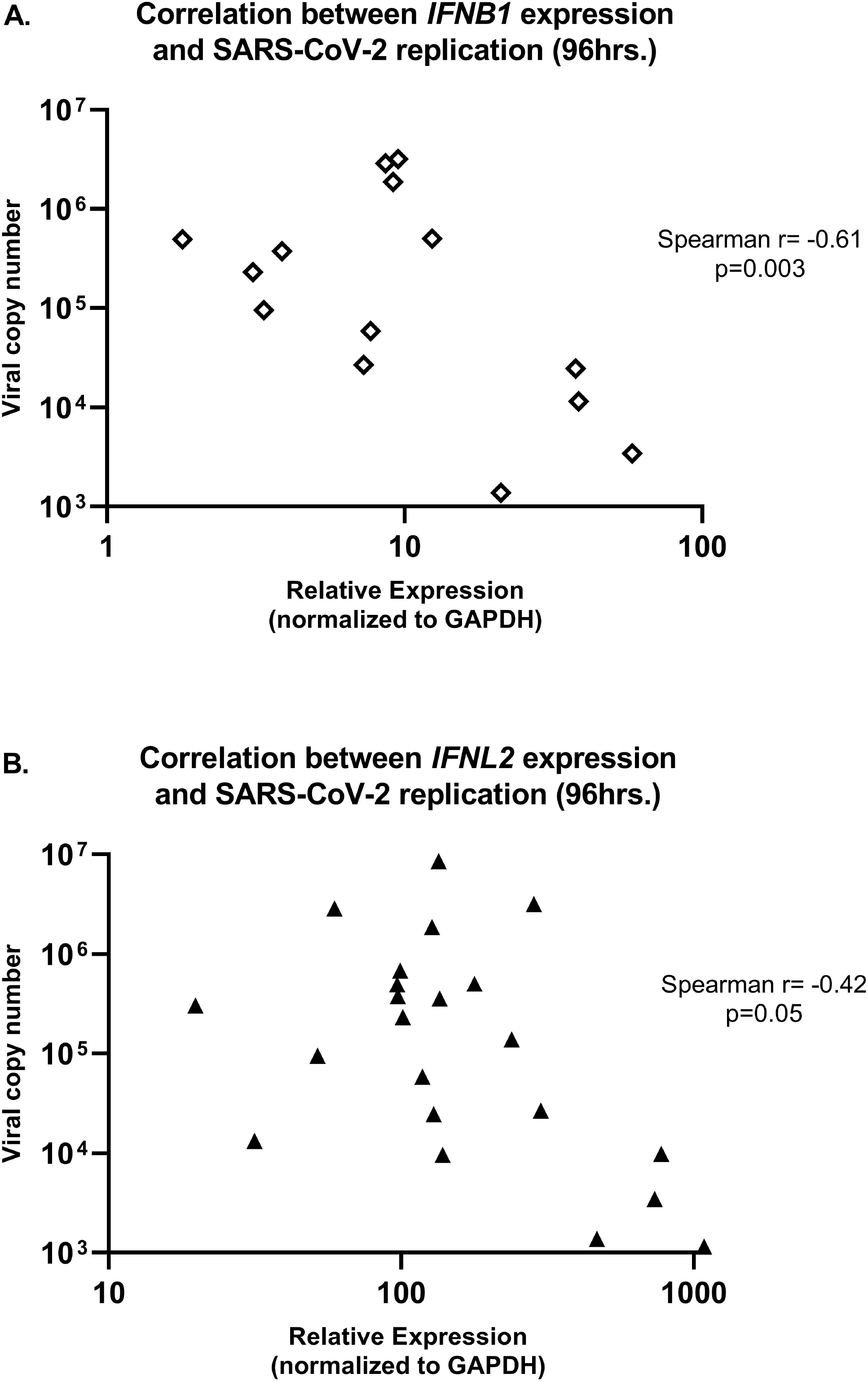
Correlation between relative gene expression of *IFNB1* or *IFNL2* (normalized to GAPDH) and SARS-CoV-2 viral replication by quantitative PCR in primary bronchial epithelial cell cultures in children (n=14) and adults (n=8). *IFNB1* and *IFNL2* gene expression were inversely correlated with SARS-CoV-2 replication (**Panel A**, Spearman r=-0.61, p=0.003; **Panel B**, Spearman r=-0.42, p=0.05).

**Figure 5.**
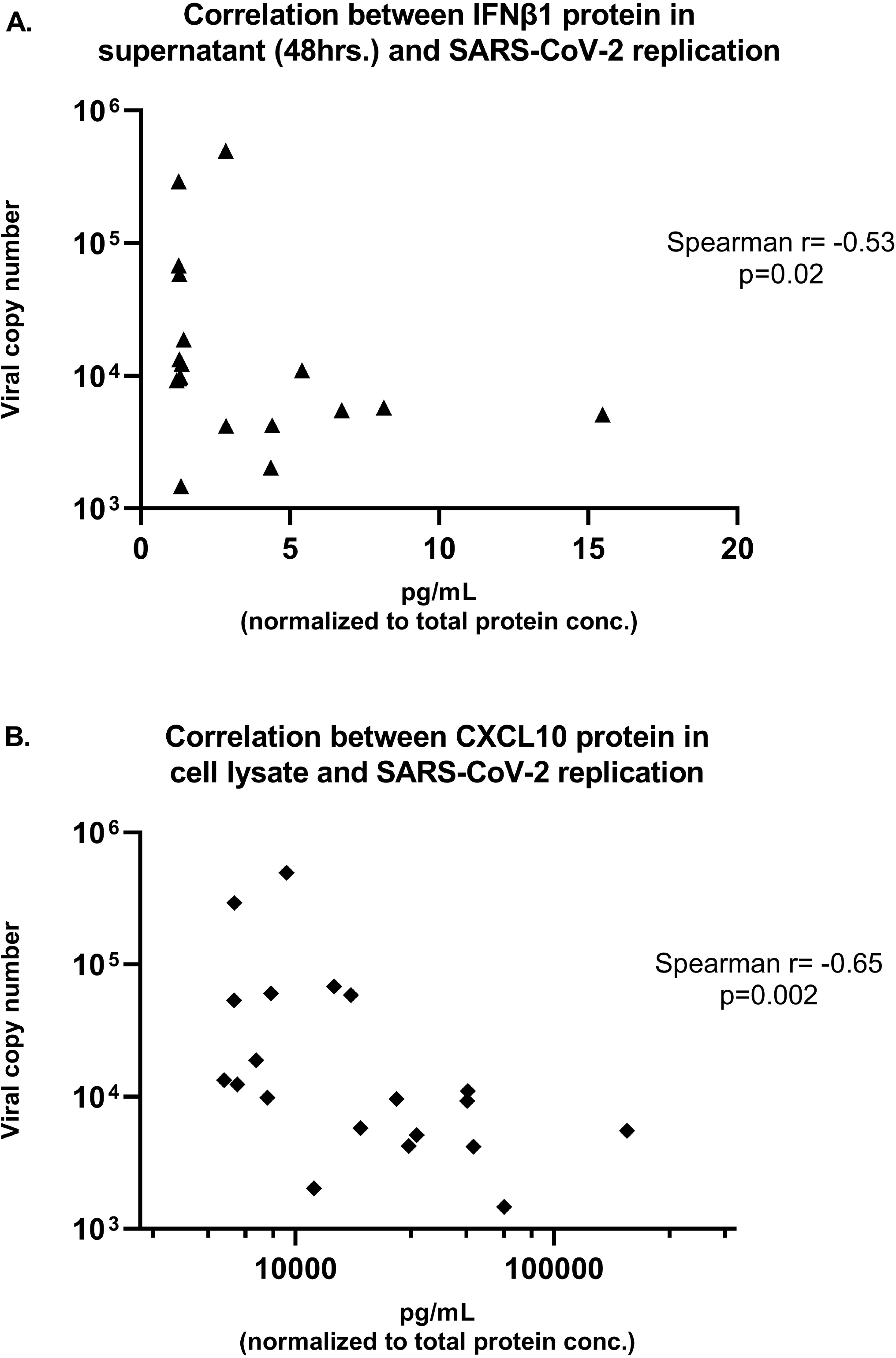
Correlation between secreted IFNβ1 concentration 48 hours after SARS-CoV-2 infection or CXCL10 concentration from the cell lysate 96 hours after SARS-CoV-2 infection (normalized to total protein concentration) and 96-hour replication of SARS-CoV-2 by quantitative PCR in primary bronchial epithelial cell cultures in children (n=14) and adults (n=8). Secreted I FNβ1 and SARS-CoV-2 replication were significantly inversely correlated (**Panel A**; Spearman r=-0.53, p=0.02), and CXCL10 concentration from cell lysates was also significantly inversely correlated with SARS-CoV-2 replication (**Panel B**; Spearman r=-0.65, p=0.002).

In organotypic bronchial epithelial cultures from 14 children and 10 older adults, replication of SARS-CoV-2 was compared between cultures infected with SARS-CoV-2 alone (MOI=0.5), infection of cultures with HRV-16 (MOI=0.5) followed 72 hours later by infection with SARS-CoV-2 (MOI=0.5), infection of I FNβ1 pre- and concurrently treated cultures with SARS-CoV-2, and infection of IFNλ2 pre- and concurrently treated cultures with SARS-CoV-2. Pre-infection of bronchial AECs with HRV-16 led to a marked reduction in SARS-CoV-2 replication 96 hours following infection (median SARS-CoV-2 copy number 267,264 vs. 14,788, p=0.002; Figure 6). Treatment of AEC cultures with recombinant IFNβ1 reduced SARS-CoV-2 replication from a median copy number of 267,264 to 11,947 (p=0.0001) and treatment of AEC cultures with recombinant IFNλ2 reduced SARS-CoV-2 replication from a median copy number of 267,264 to 11,856 (p=0.0002).

**Figure 6.**
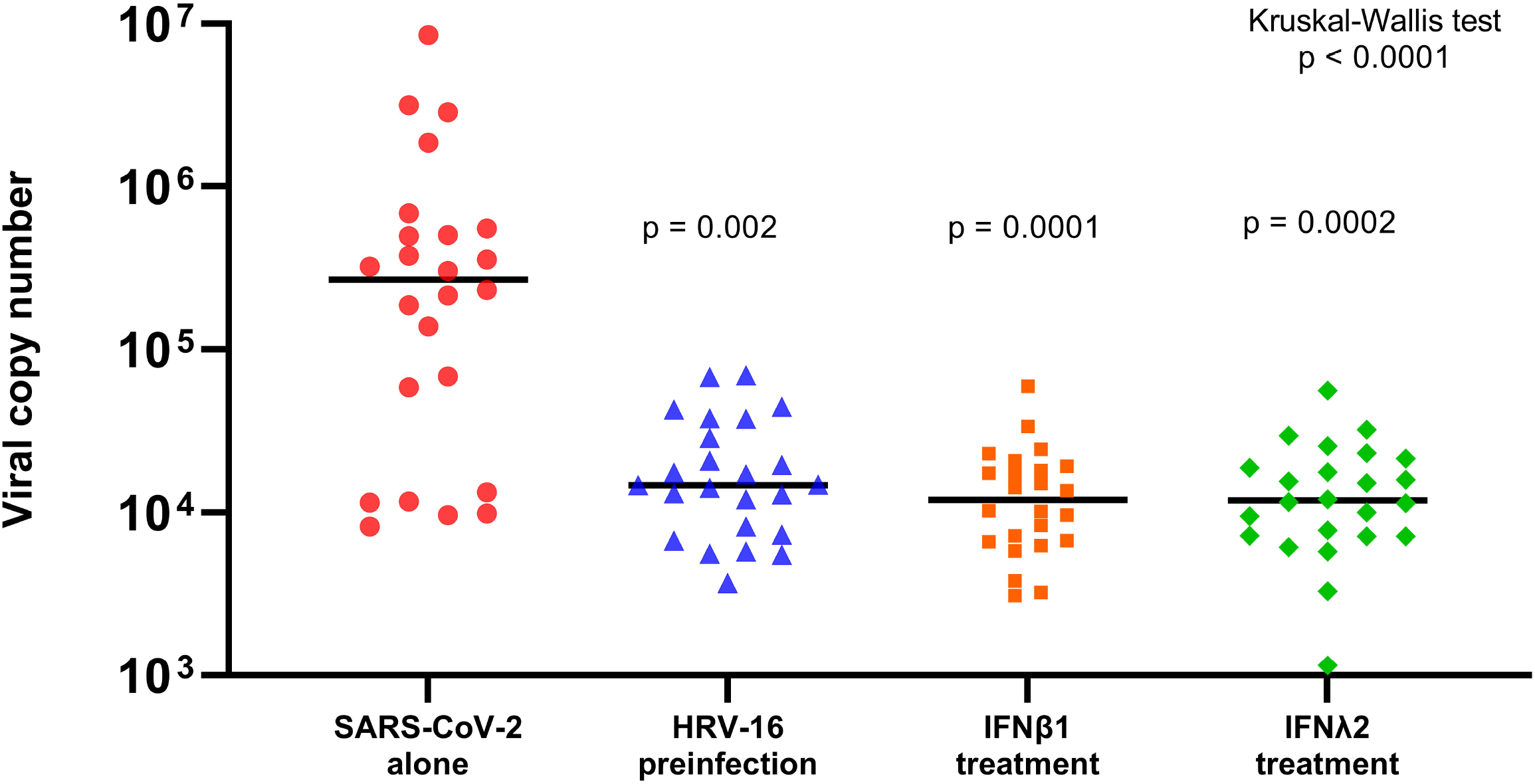
SARS-CoV-2 replication by quantitative PCR in primary bronchial airway epithelial cell cultures from children (n=14) and adults (n=10) infected in parallel with SARS-CoV-2 alone at MOI=0.5 (red circles), SARS-CoV-2 infection 72 hours following pre-infection with HRV-16 (MOI=0.5: blue triangles), pre- and concurrent treatment with recombinant IFNβ1 (orange squares), and pre- and concurrent treatment with recombinant IFNλ2 (green diamonds). Viral copy number was quantified by PCR in RNA harvested 96 hours after SARS-CoV-2 infection. Pre-infection of primary bronchial AECs with HRV-16 significantly reduced SARS-CoV-2 replication (median copy number 267,264 vs 14,788, p=0.002). Treatment of bronchial AEC cultures with recombinant IFNβ1 or IFNλ2 also significantly reduced SARS-CoV-2 replication (median copy number 267,264 to 11,947, p=0.0001; median copy number 267,264 to 11,856, p=0.0002, respectively). Kruskal–Wallis one-way ANOVA on ranks was used to compare all experimental conditions. Dunn’s test was used for comparisons between SARS-CoV-2 alone and individual experimental conditions. Bars indicate median values.

Given that SARS-CoV and MERS have been noted to evade innate antiviral defenses at various steps between viral sensing and transcription and translational of type I and III interferons, and ultimately transcription of an array antiviral genes(12) (13) (14) (15) (16) (17), we assessed one potential proximal step where SARS-CoV-2 may evade sensing of viral nucleic acids by comparing gene expression of the pattern-recognition receptor and RNA viral sensor IFIH1/MDA5 between primary bronchial AEC cultures infected in parallel with SARS-CoV-2 (MOI=0.5) or HRV-16 (MOI=0.5). We observed that IFIH1 expression was more than 2-fold greater following infection with HRV-16 as compared to following SARS-CoV-2 infection (Figure 7; p=0.003).

**Figure 7.**
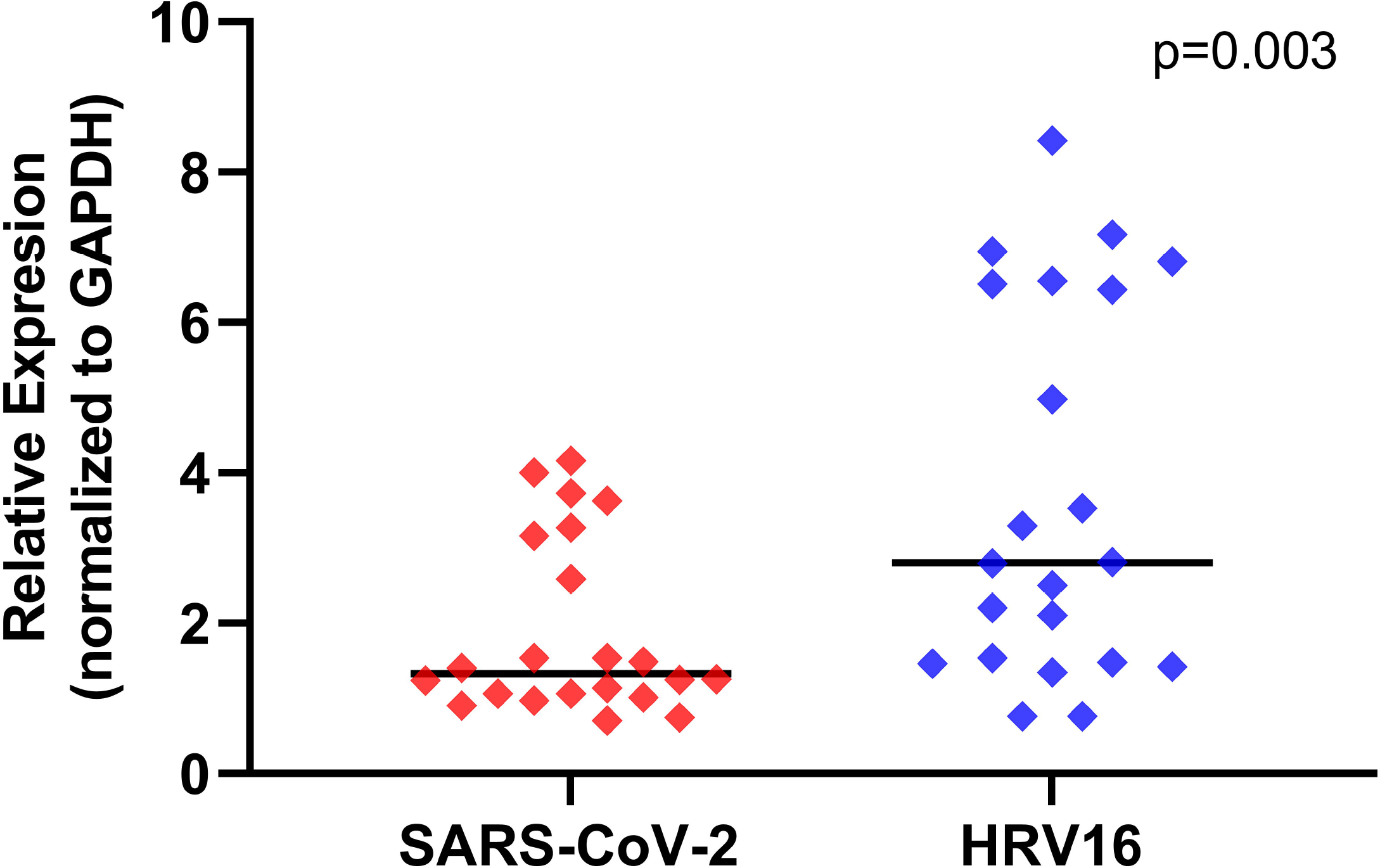
Relative gene expression of pattern-recognition receptor and RNA viral sensor *IFIH1 (MDA5)* (normalized to GAPDH expression) by primary bronchial airway epithelial cell cultures in children (n=14) and adults (n=8). Gene expression was quantified by PCR in RNA harvested 96 hours after parallel infection with SARS-CoV-2 (MOI=0.5, red diamonds) or HRV-16 (MOI=0.5, blue diamonds). IFIH1 expression was significantly higher after HRV-16 infection than after SARS-CoV-2 infection (p=0.003 by Wilcoxon matched-pairs signed rank test; bars indicate median values).

## DISCUSSION

A growing body of literature suggests that beta-HCoVs, including SARS-CoV-2 appear able to antagonize type I and III IFN responses at mucosal surfaces at multiple steps between viral sensing and production of interferon induced antiviral proteins(18) (19) (33). In this study we directly compared type I and III IFN responses to SARS-CoV-2 and HRV-16 infection by primary organotypic bronchial AEC cultures from children and adults, and assessed the impact of exogenous treatment with recombinant IFNβ1 or IFNλ2 on SARS-CoV-2 replication as well as the impact of heterologous infection with HRV-16 prior to SARS-CoV-2. We observed significant heterogeneity in SARS-CoV-2 replication between primary AEC lines from different human donors, however, despite between donor heterogeneity we also observed that SARS-CoV-2 replicated approximately 100 times more efficiently than HRV-16 in these primary bronchial AEC cultures. As compared to uninfected cultures, the relative increase in expression of *IFNB1, INFL2, and CXCL10* following infection with HRV-16 was significantly greater than following infection with SARS-CoV-2, and the protein concentrations of type I and III IFN and the IFN stimulated chemokine CXCL10 in both cell lysates and supernatant were significantly greater in AEC cultures following infection with HRV-16 as compared to SARS-CoV-2. In SARS-CoV-2 infected AEC cultures type I and III IFN gene expression and protein production were inversely correlated with viral replication. Furthermore, treatment of AEC cultures with recombinant IFNβ1 or IFNλ2, or pre-infection of AEC cultures with HRV-16, markedly reduced SARS-CoV-2 replication.

Sensing of beta-HCoVs by the innate immune system is believed to be primarily through pattern recognition receptors (PRRs), including cell surface or endosomal transmembrane TLRs TLR3 and TLR7, the cytosolic RIG-I-like receptors melanoma differentiation-associated protein 5 (MDA5), as well as retinoic acid-inducible gene I (RIG-I) (6) (7) (8) (9). PRR’s then mediate activation of signaling cascades leading to induction of type I and III IFN responses(6) (7) (8) (9). Recently Sampaio et al. reported that in the lung cancer cell line Calu-3 the cytosolic RNA sensor MDA5 was required for type I and III IFN induction when cells were infected with SARS-CoV-2 infection(6).

Studies using immortalized cell lines (e.g. Vero, HeLa, Calu-3, 293T) *in vitro,* as well as murine *in vivo* studies, have suggested a number of potential mechanisms by which beta-HCoVs (e.g. SARS-CoV, MERS-CoV, and SARS-CoV-2) may evade IFN responses at the level of the airway epithelium. Prior to the onset of the COVID-19 pandemic, these mechanisms were investigated extensively for SARS-CoV and MERS-CoV. One group of beta-HCoV proteins, the predominantly non-structural proteins (nsps), are recognized to have IFN-antagonistic impacts. Several nsps (e.g. nsp1 and nsp3) interfere with signal transduction mediated by PRRs, while other nsps evade recognition by PRRs in mucosal epithelial cells by modifying features of the viral RNA(34). There is growing evidence that SARS-CoV-2, much like SARS-CoV and MERS-CoV, has evolved a number of immune evasion strategies that may interfere with PRR’s themselves(35) (36) (37) (38) (39) (40) (41) (42), inhibit multiple steps in the signaling cascade leading to induction and translation of type I and III IFNs(43) (44) (45) (46) (47) (48) (49) (50) (51) (52), and interfere with the actions of IFNs by impeding the signaling pathways that lead to transcription and translation of anti-viral interferon stimulated genes (ISGs) (18) (53) (54) (55) (56). Lei et al. demonstrated that the SARS-CoV-2 proteins NSP1, NSP3, NSP12, NSP13, NSP14, ORF3, ORF6 and M protein all have some ability to inhibit Sendai virus-induced IFN-β promoter activation, and that ORF6 has inhibitory effects on both type I IFN production as well as signaling downstream of IFN-β production(18). Early in the COVID-19 pandemic Blanco-Melo et al. reported results from a transcriptome profiling study of various immortalized cell lines which demonstrated that SARS-CoV-2 infection elicited very low type I and III IFN and limited ISG responses, while inducing expression of pro-inflammatory cytokines genes(19), raising the possibility that a deficient epithelial IFN response to SARS-CoV-2 may facilitate enhanced local viral replication that ultimately might lead to a dysregulated systemic pro-inflammatory response.

Data from several clinical studies have provided additional support for the hypothesis that a muted initial local IFN response to SARS-CoV-2 in the airway epithelium, at least in some hosts, allows the virus to replicate unimpeded which then sets up the host for potential systemic inflammatory responses that contribute to COVID-19 pathology and severity(19) (22) (23) (24). Recently, Ziegler et al. published transcriptomics results from nasopharyngeal swabs from 15 healthy adults, 14 adults with mild COVID-19 and 21 adults with severe COVID-19, and observed that nasal epithelial cells from patients with severe COVID-19 exhibited less robust expression of anti-viral IFN response genes as compared to patients with mild COVID-19 and healthy adults, supporting their conclusion that a “failed” nasal epithelial innate anti-viral response may be a risk factor for severe COVID-19(25).

In the early stages of the pandemic, morbidity and mortality was skewed toward older patients with significant underlying comorbidities, however, over time it has become increasingly clear that clinical outcomes with COVID-19 following infection with SARS-CoV-2 is heterogeneous with outcomes even in young adults and children without medical comorbidities unpredictably ranging from ranging from asymptomatic infection to death(57). An objective of our study was to determine if heterogeneity in airway epithelial IFN responses to SARS-CoV-2 between individual pediatric and adult donors was associated with SARS-CoV-2 replication. A striking observation in our data is the marked between-donor heterogeneity in the replication of SARS-CoV-2 in organotypic AEC cultures using standardized protocols and uniform viral inoculation doses. A potential important future area of investigation will be to investigate possible genetic and epigenetic factors that may partially explain heterogeneity in SARS-CoV-2 replication in airway epithelium.

Our group and others have demonstrated that the SARS-CoV-2 entry receptor ACE2 is an ISG(20) (21). We have demonstrated that HRV-16 infection induces a type I and III interferon response, and increases ACE2 expression(21), leading us to originally speculate that HRV pre-infection of AECs might increase replication of SARS-CoV-2 through greater expression of the entry receptor and be a clinical risk factor for acquisition of COVID-19. However, our results in this study demonstrate that even through the SARS-CoV-2 entry factor ACE2 is an ISG, HRV-16 infection induces a much more potent type I and III IFN responses than SARS-CoV-2 and that heterologous infection of organotypic AEC cultures with HRV-16 three days prior to inoculation with SARS-CoV-2 markedly reduces replication of SARS-CoV-2. This suppression of SARS-CoV-2 replication was similar to the effects of exogenous treatment with IFNβ1 or IFNλ2, suggesting that the pronounced induction of these genes by HRV-16 was responsible for these findings. These findings extend upon several other recent reports including Cheemarla et al. who reported experiments in differentiated primary airway epithelial cultures from small number of adult donors and observed that infection with HRV-01A prior to infection with SARS-CoV-2 accelerated induction of ISGs and reduced SARS-CoV-2 replication(58). Similarly, Dee et al. used differentiated primary airway epithelial cultures from a single human donor, to characterize viral replication kinetics of SARS-CoV-2 with and without co-infection with rhinovirus and observed that pre-infection with HRV-16A reduced SARS-CoV-2 replication(59). Neither of these prior studies included primary ALI cultures from a robust sample size of adults and children or compared interferon and ISG responses between parallel HRV and SARS-CoV-2 infections in addition to HRV pre-infection to determine if this phenomenon is consistent across donors with heterogenous interferon responses to both viruses. Although difficult to definitely test, given that the incidence of COVID-19 in children was very low in the early months of the pandemic, during the peak of the late winter viral respiratory season in the United States and Europe when rhinovirus, RSV, and influenza activity was high, our data together with the studies by Dee and Cheemarla (58, 59) lead us to hypothesize that high rates of typical respiratory viral pathogens among Children in the northern hemisphere in February-March 2020 may have contributed to protection of children early in the COVID-19 pandemic by generally inducing airway type I and III IFN responses.

Given the steady evolution of new SARS-CoV-2 variants through 2021 and continued significant resistance to vaccination among a sizable minority of people with access to vaccines, the pandemic has continued to result in high levels of morbidity and mortality in many areas of the world, fueling an ongoing need for therapeutics to treat COVID-19. Our results demonstrating marked reduction in SARS-CoV-2 replication in AEC cultures treated with recombinant IFNβ1 or IFNλ2 provides further mechanistic evidence to support the possible use of inhaled interferon as a possible treatment option if initiated early enough during COVID-19. A recent randomized, double-blind, placebo-controlled, phase 2 trial of inhaled nebulized interferon beta-1a (SNG001) for treatment of SARS-CoV-2 infection demonstrated that patients who received SNG001 early in their disease course had greater odds of improvement and recovered more rapidly from SARS-CoV-2 infection than patients who received placebo, providing a strong rationale for further trials of this agent(60).

We are not aware of other studies to date that have directly compared innate immune responses between SARS-CoV-2 and HRV in organotypic AEC cultures from many pediatric and adult donors. However, there are several limitations of our primary airway epithelial model system. First, our *ex vivo* system lacks interaction with immune cells and the complex immune responses that occur in vivo in the context of COVID-19, and therefore we cannot assess how heterogeneity in interferon responses to SARS-CoV-2 at the level of the airway epithelium relate to systemic immune responses or clinical outcomes *in vivo.* Second, in this study we did not investigate potential genetic or epigenetic factors that may explain the between subject heterogeneity in interferon responses and viral replication that we observed. Finally, given the limitations posed by the complex logistics of completing these experiments in a biosafety level 3 (BSL-3) facility, together with limitations in available material from organotypic cultures from a sizeable number of human donors, we were constrained in the number of feasible sample harvesting timepoints which prevented us from conducting a high resolution assessment of the time kinetics of viral infection and interferon responses in the present study; however, our choice to harvest supernatant 48 hours following SARS-CoV-2 infection and RNA 96 hours following infection was informed by both our prior work with RSV(29) and preliminary experiments with SARS-CoV-2 (data not shown) where we observed that in organotypic primary ALI cultures type I and III interferon responses peak between 24-48 hours while expression of downstream ISGs peak between 72-96 hours.

In conclusion, in this study we have demonstrated that in addition to remarkable between subject heterogeneity in interferon responses and viral replication, SARS-CoV-2 elicits a less robust type I and III interferon response in organotypic primary bronchial AEC cultures than does human rhinovirus, and that pre-infection of AECs with HRV-16, or pre-treatment with recombinant IFN-β1 or IFN-λ2, markedly reduces SARS-CoV-2 replication.

## List of Abbreviations

COVID-19: coronavirus disease 2019
IFN: interferon
PRR: pattern recognition receptors
MDA5: melanoma differentiation-associated protein 5
RIG-I: retinoic acid-inducible gene I
TLR: toll-like receptor
HCoV: human coronavirus
HRV-16: human rhinovirus-A16
AEC: airway epithelial cell
ALI: air-liquid interface
BSL-3: Biosafety Level 3
MOI: multiplicity of infection
qPCR: quantitative polymerase chain reaction

## Declarations

### Ethics Approval

Airway epithelial cells from children were obtained under study #12490 approved by the Seattle Children’s Hospital IRB. Parents of subjects provided written consent and children over 7 years of age provided assent. Airway epithelial cells from adults were purchased from Lonza® without personal identifiers. The Seattle Children’s Hospital IRB determined that use of de-identified adult airway epithelial cells purchased from Lonza® did not require ethics approval or consent.

### Consent for publication

This manuscript does not contain any individual person’s data in any form.

### Availability of data and materials

The datasets used and/or analysed during the current study are available from the corresponding author on reasonable request.

### Author Contributions

Conceptualization, K.A.B., C.T., J.S.D.; methodology, K.A.B, E.R.V., L.M.R., O.O., J.S.D.; validation, E.R.V., L.M.R., K.A.B., J.S.D.; formal analysis, E.R.V., J.S.D.; investigation, E.R.V, L.M.R, K.A.B., M.P.W., O.O., J.S.D.; resources, T.S.H., J.S.D.; data curation, O.O., K.A.B., L.M.R., M.P.W., E.R.V., J.S.D.; writing—original draft preparation, E.R.V, J.S.D.; writing—review and editing, E.R.V., L.M.R., M.P.W., O.O., S.F.Z., T.S.H., D.F.R., C.T., K.A.B., J.S.D.; supervision, J.S.D.; project administration, J.S.D.; funding acquisition, S.F.Z, J.S.D. All authors have read and agreed to the published version of the manuscript.

